# Chronic silencing of corticothalamic layer 6 pyramidal cells affects cortical excitability and tactile behavior

**DOI:** 10.1101/215558

**Authors:** Desire Humanes-Valera, Jordi Aguila, Veronika Gondzik, Karen A. Moxon, Martin K. Schwarz, Patrik Krieger

## Abstract

Cortical projections originating in layer 6 (L6) in mouse primary somatosensory cortex have an important function controlling both cortical and sub-cortical activity. To study cortical plasticity and communication between somatosensory and motor cortex, L6-Ntsr1 cells were chronically silenced using tetanus toxin and the effect this had on cortical activity and behavior was investigated. A 2 Hz stimulation protocol was used to potentiate whisker evoked local field potentials (LFP) in a layer-dependent manner in both cortices. Silencing L6 pyramidal cells, the LFP potentiation in both cortices was affected without a change in the spontaneous multi-unit activity. Animals with L6 chronically silenced used more low-amplitude whisks, which presumably compensates for a reduction in cortical excitability. These results suggest that L6 is not only an important cortical output layer that modulates sub-cortical circuits, but also that it controls cortical dynamics.

## Introduction

The acquisition of sensory information is an active process where sensory input and motor behavior influence each other (Gibson, 1962; Wolpert et al., 1995; Ahissar and Arieli, 2001; Kleinfeld et al., 2002; Nelson and MacIver, 2006; Diamond et al., 2008); a good model to study sensorimotor integration is the vibrissal system in rodents (Krieger and Groh, 2015), the function of which can be compared with digital palpation and saccadic eye movements in primates (Carvell and Simons, 1990; Diamond, 2010). Rodents explore the environment moving their whiskers at different frequencies to identify different objects and surfaces (Carvell and Simons, 1990; Wolfe et al., 2008). Presumably, the changes in whisking frequency not only increase the sampling of touch information, but also the increased sensory sampling frequency triggers cortical plasticity related to learning. In sensory systems, peripheral stimulation to different frequencies has been shown to produce long-term potentiation of cortical responses in sensory cortices in both humans and rodents (Clapp et al., 2005; Teyler et al., 2005; An et al., 2012). Furthermore, cortical plasticity caused by peripheral sensory stimulation can extend to a potentiation also in motor cortex (Megevand et al., 2009). Cortical potentiation is affected by activity in the cortico-thalamo-cortical loops. Important for regulating activity in these pathways are layer 6 (L6) pyramidal cells which can modulate both thalamic response properties and cortical activity (Olsen et al., 2012; Kim et al., 2014; Mease et al., 2014; Velez-Fort et al., 2014; Crandall et al., 2015; Denman and Contreras, 2015; Hirai et al., 2017). Little is known about the role of L6 cells in sensorimotor integration, a process that involve communication between sensory and motor areas.

The aim of the present investigation was to examine the importance of somatosensory layer 6 in controlling whisker-evoked cortical activity in somatosensory and motor cortex. Furthermore, the effect on a whisker-dependent behavior was studied. For this purpose, we used transgenic mice (Ntsr1-cre) with a population of genetically defined layer 6 pyramidal cells (Gong et al., 2007). Using the cre-loxP recombination system, expression of toxins was targeted to these cells. To silence synaptic transmission, we used tetanus toxin to induce a permanent block. Silencing the activity of these cells, we investigated the importance of L6-Ntsr1 activity for plasticity and whisker-dependent behavior.

## Materials and methods

All experiments were in accordance with the local government ethics committee (Landesamt für Natur, Umwelt und Verbraucherschutz, Nordrhein-Westfalen). Two separate silicon microelectrode arrays were used for extracellular recordings simultaneously in somatosensory cortex and motor cortex in ten WT mice (7 male, 3 female) and ten Ntsr1-cre (GENSAT founder line GN220) mice (4 male, 6 female). Behavioural testing in the gap-crossing task was performed in seven WT mice (3 male, 4 female) and ten Ntrs1-Cre mice (4 male, 6 female). All animals were littermates and housed together until experiments were performed.

### Stereotaxic Viral Injections

Stereotaxic injections were performed in male and female WT and Ntsr1-cre mice between 3 and 5 months old. Animals were anesthetized with xylazine (16 mg/kg) / ketamine (97 mg/kg). The body temperature was kept constant (37 °C) using an automatically controlled heating pad (FHC, ME, USA). Animals were placed in a stereotaxic frame (Model 1900; David Kopf Instruments, CA, USA). The skin of the head was softly removed and the skull was exposed. For the electrophysiology experiment we performed a craniotomy on the left hemisphere over the vibrissal primary somatosensory cortex (SCx): antero-posterior (to bregma) from -1 to -2mm; Lateral (to midline) from 3 to 4mm (Franklin and Paxinos, 2008). Then 1400 nl of Adenoassociated viral (AAV) particles encoding for rAAV-EF-doublefloxed-TetTX-ko (in Ntsr1-Cre animals) or encoding GFP (in WT animals), was distributed equally over 7 target sites in the somatosensory cortex (200 nl in each injection place) with the following coordinates [in mm] relative to bregma, midline, and dura respectively: (1.) -2, 3, 0.9; (2.) - 2, 3.5, 0.9; (3.) -2, 4, 0.9; (4.) -1, 3, 0.9; (5.) -1, 3.5, 0.9; (6.) -1, 4, 0.9; and (7.) -1.5, 3.5, 0.9. For the gap-crossing task, a craniotomy was made on the left hemisphere over the vibrissal primary somatosensory cortex (SCx) with the following coordinate: -1.7mm antero-posterior; 3.1mm lateral (to midline) and 0.9 mm of depth. In the target site 500 nl of adenoassociated viral (AAV) particles encoding for rAAV-EF-doublefloxed-TetTX-KO was injected in animals used for gap-crossing task. Subsequently, mice were sutured and housed in their cages until the experiment. Incubation time was 3 weeks. Ntsr1-cre mice injected with tetanus toxin are referred to as “L6-Silenced animals”.

### Experimental protocol

Animals were anesthetized with urethane (1.5g/kg i.p.). The body temperature was kept constant (37 °C) using an automatically controlled heating pad (FHC, ME, USA). Animals were placed in a stereotaxic frame (SR-6M-HT; Narishige Scientific Instruments, Japan). The skin of the head was softly removed and the craniotomy over vibrissal primary somatosensory cortex (SCx) was exposed. A second craniotomy was performed on the left hemisphere over the vibrissal primary motor cortex (MCx; coordinates: lateral 1mm, antero-posterior 1mm). Once the electrodes were positioned at the corresponding area and correct depth in SCx and MCx respectively (see section “extracellular recordings”, below), spontaneous activity and evoked responses were recorded in “control condition” (0.2 Hz stimulation). After the control period, the whiskers were stimulated using the potentiation protocol (2 Hz, during 10 min). Thereafter, we stimulated at 0.2 Hz at different times to measure the LFP response over time (10, 25, 45 min and 1 hour after the 2 Hz protocol, Fig. 1A). After the experiment, each mouse was transcardially perfused with phosphate-buffered saline (PBS) followed by 4% paraformaldehyde in PBS before the brain was removed. Brains were saturated overnight in 4% paraformaldehyde and then coronally transected in 50 µm slices. In WT animal we used Neurotracer (Neuro Trace Fluorescent Nissl Stains (N-21480; Invitrogen), Fig. 1B,C) to visualize neuron cell bodies in every layer, and based on cell densities determine layer borders in SCx and MCx slices. In L6-Silenced animals, we in addition made immunohistochemistry to identify the tetanus-toxin transfected neurons.

**Figure 1.**
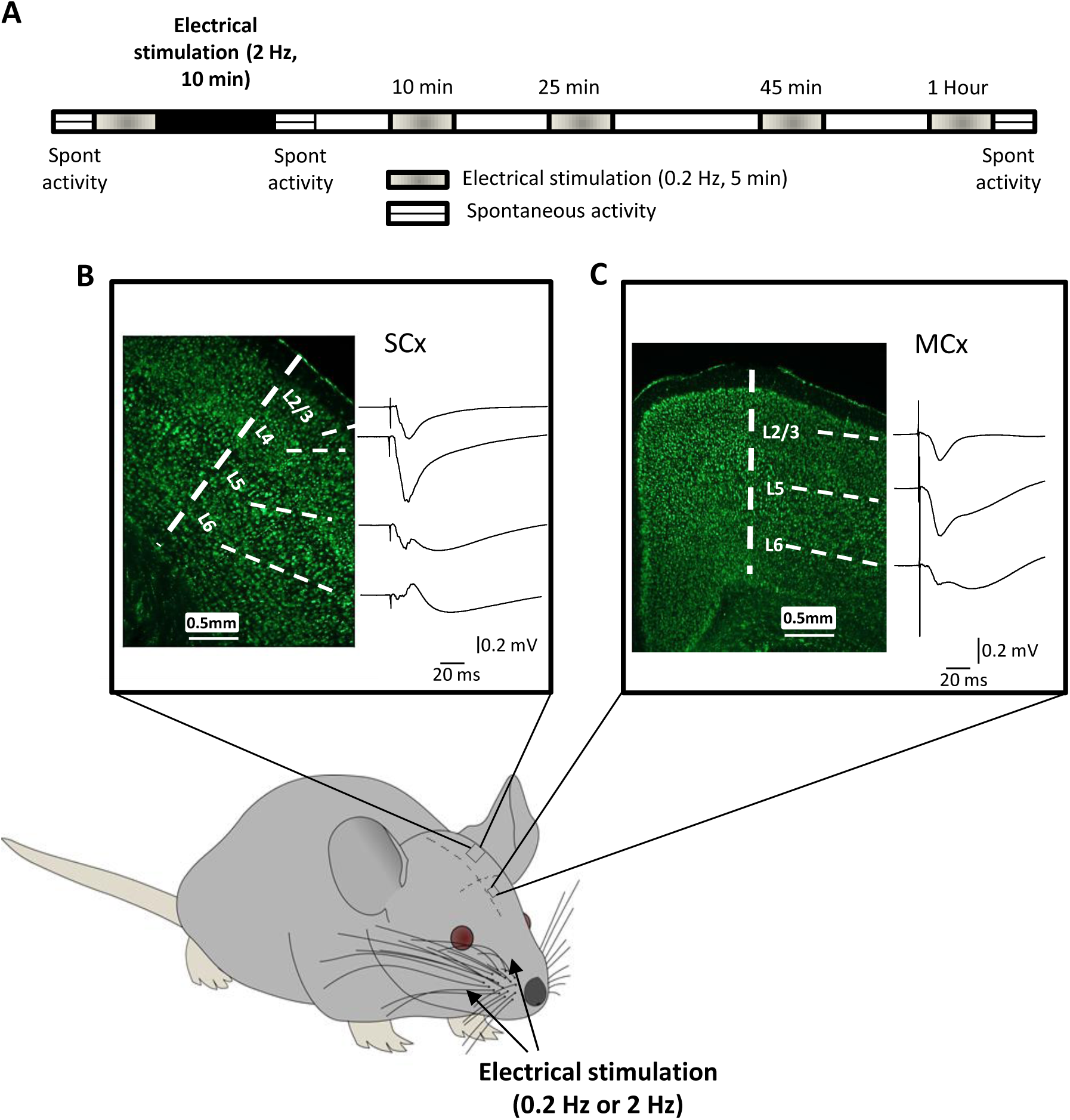
Extracellular recordings from different layers in somatosensory and motor cortex. **(A)** Experimental protocol. Spontaneous activity and whisker evoked responses were measured at different time points. Whisker deflections were induced by electrical stimuli delivered to the whisker pad at two frequencies, 0.2 Hz (test stimulation) and 2 Hz (potentiation protocol). The whisker evoked LFP amplitude was measured during a 5 min period before and at different time points after the 2 Hz potentiation (10, 25, 45 and 60 min). **(B) and (C)** Extracellular recordings with 32-Channel silicon probes were made from vibrissal primary somatosensory cortex (SCx) and vibrissal primary motor cortex (MCx) in urethane anaesthetized mice. **(B)** Coronal section of somatosensory cortex with Neurotracer staining (green) and the average LFP response characteristic for each layer and **(C)** coronal section of motor cortex with Neurotracer staining and average LFP responses in each layer. Neurotracer was used to visualize the cell bodies and the white line shows the electrode trajectory.

### Immunohistochemistry

50 µm thick brain slices were submerged in PBS, washed 3 times for 10 min in PBS plus 0.2% Triton (T-PBS). The sections were permeabilized for 30 min in 0.25% T-PBS followed by blockade for 1 hour in T-PBS plus 20% Goat-Serum (art. no. 005-000-121; Dianova, Germany). We diluted the antibody for the fluorescent tag on the tetanus toxin AAV Kusabira–Orange (Anti-monomeric Kusabira-Orange 2 pAb (MBL-PM051); Amalgaan, Japan), and NeuN (art. no. MAB377; Merck Millipore) in 0.2% T-PBS plus 10% Goat-Serum, 1:500 and 1:100 respectively and then the sections were incubated overnight in this mix. The next day the sections were washed 3 times for 10 min in T-PBS. After washing slices were incubated with second antibodies diluted in T-PBS plus 1% Goat-Serum for 90 min in room temperature. As second antibodies we used Goat-Anti-Rabbit-Cy3 (diluted 1:500; art. no. 111-165-033; Dianova, Germany) to detect Kusabira-Orange (i.e., tetanus toxin infected cells), and Goat-Anti-Mouse-Cy2 (diluted 1:250; art. no. 115-225-166, Dianova, Germany) for detecting NeuN labeled cells. Finally, slices were washed 3 times for 10min in PBS, and then mounted using MOWIOL 4–88 (SIGMA).

### Whisker pad stimulation

For extracellular recordings with silicon probes, electrical pulses (0.5ms pulse duration) were applied to the whisker pad using bipolar needle electrodes located subcutaneously at sites anterior and posterior (Farkas et al., 1999; Castro-Alamancos, 2004; Moxon et al., 2007; Chakrabarti et al., 2008). The rationale for using electrical stimulation was to maximize the area activated within the whisker pad (Chakrabarti et al., 2008). This type of stimulation evokes neuronal responses that may be caused by both movements of the whiskers and direct stimulation of the sensory nerve. Two stimulation protocols were applied (Fig. 1): 0.2 Hz during 5 min as background stimulation, and 2 Hz during 10 min as potentiation protocol. Current pulse amplitude was adjusted to levels (15-20 µA) that evoked movement of the whisker pad. In two-thirds of the experiments the whisker movements were synchronously tracked using a high speed camera (300 frames per seconds; Pike F-032B, ALLIED Vision Technologies GmbH, Germany). The electrical stimulation moved on average 8.0 ± 0.7 whiskers, and the average angle of this movement was 10.5 ± 1.3 degrees.

### Electrophysiological recordings

Extracellular recordings were obtained with two 32 channels silicon electrode arrays (A1x32-6mm-50-177; NeuroNexus Technologies, MI, USA) in combination with a Multichannel Acquisition Processor (MAP, Plexon Inc., TX, USA). One silicon probe was stereotaxically lowered into vibrissal primary somatosensory cortex (Lateral, 3.5 mm; Antero-posterior, 1.5 mm) and another one into vibrissal primary motor cortex (Lateral, 1mm; Antero-posterior, 1 mm). The trajectory of the probes (dotted line in Fig. 1) was confirmed using DiI labeling (D282; Life Technologies GmbH, Germany). The layer position of each electrode was determined based on the electrode tract and the layer borders. Layer borders were determined from the changes in cell body density in the different layers (Groh et al., 2010). Cell bodies (green in Fig. 1B,C) were labeled with Neurotracer (Neuro Trace Fluorescent Nissl Stains (N-21480; Life Technologies GmbH, Germany)). The distance between channels is 50 µm, thus to ensure a correct layer mapping of each channel, one to four channels were omitted from each layer. The average LFP response for a given layer was averaged across 3-5 electrodes.

## Behavioural test

Whisker kinematics and behavior were studied in a gap-crossing apparatus built as previously described (Juczewski et al., 2016). Experiments were done using seven WT mice and ten L6-Silenced mice. The animals were habituated to the experimenter and to the gap-crossing apparatus 2 days prior to a behavioral test. The habituation included 20 min inside the apparatus with the platforms pushed together, so that the animals could cross between the platforms without a gap between them. On the first day, the animal was placed inside the apparatus with background white noise and lights on; on the second day, background white noise but now with lights turned off. After the second habituation session, all whiskers, except C2 on right side of the snout, were removed. The whisker removal procedure was done under isoflurane anesthesia. The removed whiskers were trimmed with scissors to fur-level or plucked following the daily test session. Trimming was done after the daily testing to avoid stress during the task. The mice are able to learn the gap-crossing task with a single whisker (Celikel and Sakmann, 2007).

The movement of the mouse within the behavioral apparatus was monitored with infrared motion sensors (MS). Motion sensors close to the gap on either side of the platform are called MS2 and MS3 respectively. An “attempt” is defined as an event where the animal activates (by breaking the beam) the MS close to the gap (MS2 or MS3) and a “successful attempt” is an event where the animal actually crosses over the gap to reach the other platform. Exploration duration for a successful attempt was defined as M2 ON, M2 OFF, M3 ON, and as MS3 ON, MS3 OFF, MS2 ON. The ON and OFF time of the beam breaks from each MS were analyzed in MATLAB using custom-written routines. In essence, these parameters will quantify for how long time the animal explores the gap.

Testing consisted of one 20-minute session per day for 8 consecutive days. Animals were placed inside the apparatus with white noise in the background and in complete darkness. They were allowed to freely explore and cross the gap spontaneously. The gap distance was changed in increments of 0.5 cm after each successful cross according to a pseudorandom protocol that weighted larger distances as the animal made more crossings. The protocol was divided into 4 blocks. Within each block, distances were selected randomly from a predetermined range unique to that block, and the number of successful crossings that were needed before proceeding to the next block was 1 or 3. The protocol was as follows: Day 1: block 1 (3 crossings before switching to the next block), gap distance 4–4.5 cm; block 2 (3 crossings), 4.5–5.5 cm; block 3, (3 crossings), 5–6.5 cm; block 4, 5.5–7 cm; Day 2–4: same as Day 1 expect that there was only 1 successful crossing necessary in block 1, before switching to block 2; Day 5–8: block 1, (1 crossing), 4–4.5 cm; block 2, (3 crossings), 5.5–6 cm; block 3, (3 crossings), 6–7 cm; block 4, 6–7.5 cm. This pseudo-random protocol allowed mice to work up to the greater distances while maintaining a degree of unpredictability. The exact distances within these ranges varied for each mouse and each session. After each session, the animal was placed back in its home cage and the test apparatus was cleaned with 70 % ethanol.

### Data analysis

Extracellular recordings with silicon probes were made from both somatosensory and motor cortex by: (1) bandpass filtering the raw signals at high frequencies (300–3000 Hz), (2) detecting all spikes that exceeded the background noise by at least 5 SDs (Rey et al., 2015), and (3) averaging the spikes across stimuli to construct peristimulus time histograms (PSTHs). The magnitude of multiunit responses was calculated for a “short latency” response as the average number of spikes per stimulus produced in the first 50 ms after the stimulus, and for a “long latency” response as the average number of spikes per stimulus during 150-500 ms after stimulus onset. LFP responses were obtained by averaging the raw signals across 60 stimuli. LFP response amplitude was calculated as the absolute value of the negative peak in the average response. Cortical spontaneous activity was studied in recordings at least 180 s long, performed at three time points during the experimental protocol (Fig. 1A), and analyzed as multiunit responses to quantify the firing rate in Hz. All results were obtained by routines written in Matlab (The Mathworks, USA) and using tools from Neuroexplorer (Nex Technologies, AL, USA).

Gap-crossing task. Whisker tracking was done using BIOTACT Whisker Tracking Tool (Perkon et al., 2011; Zuo et al., 2011). “Successful attempts” was defined as the animal crossing over the gap to reach the opposite platform. The quantification was based on the whisking behavior data from the successful attempts only. “Duration of touch event” was calculated as the average duration for a single whisker touch, i.e., the time during which the whisker was in physical contact with the platform. Whisker touches were counted by a human observer.

### Statistical analyses

Data followed a Gaussian distribution (d’Agostino and Pearson omnibus normality test) and were tested using parametric statistics.

Extracellular recordings. Differences in multiunit responses were identified with two-way ANOVA: LAYERS with four levels in the case of SCx and three levels in MCx (L2/3, L4, L5, L6 and L2/3, L5, L6 respectively) and WT/L6-SILENCE with two levels (WT and L6-Silenced animals). Changes in the short latency responses were evaluated with a two-way ANOVA: TIME with three levels (Control condition, 45 min and 1h after potentiation stimulation), and LAYERS with four levels in the case of SCx and three levels in MCx (L2/3, L4, L5, L6 and L2/3, L5, L6 respectively). Changes in the long latency response was quantified (with a 350 ms time window) by comparing the relative change of the number of spikes during spontaneous activity and during the long-latency response, and tested with unpaired t-test.

Difference in cortical spontaneous activity were evaluated with ordinary two-way ANOVA, with SCX-MCX as one factor with two levels (SCx and MCx) and WT/L6-SILENCE as the second factor with two levels (WT and L6-silenced animals). Evaluation of the spontaneous activity in time was made with two-way ANOVA: LAYERS with four levels in the case of SCx and three levels in MCx (L2/3, L4, L5, L6 and L2/3, L5, L6 respectively) and TIME with three levels (Control condition, after potentiation protocol and the end of recording protocol).

Differences in the LFP responses between layers in control condition were identified with a one-way repeated measure (rm) ANOVA, LAYERS as factor with 4 levels in SCx (L2/3, L4, L5 and L6) and 3 levels in MCx (L2/3, L5 and L6). Evaluation of the LFP responses between WT and L6-Silenced animals in control condition was made with two-way repeated measure (rm) ANOVA, LAYERS as factor with 4 levels in SCx (L2/3, L4, L5 and L6) and 3 levels in MCx (L2/3, L5 and L6) and WT/L6-SILENCE with two levels (WT and L6-Silenced animals). Changes in LFP responses between layers at the different recording time points were evaluated with two-way repeated measure (rm) ANOVA, with repeated measure on both main factors: LAYERS with four levels in the case of SCx and three levels in MCx (L2/3, L4, L5, L6 and L2/3, L5, L6 respectively), and TIME with five levels (Control condition, 10 min, 25 min, 45 min and 1h after potentiation stimulation).

As post hoc tests we used Dunnett’s test when multiple comparison test was made with control condition, Tukey’s when we used multiple comparisons (not only with control condition) and Bonferroni test when comparison was made between two conditions. Statistical analysis was done with Prism 6 (GraphPad Software, CA, USA). The behavioral data was tested with unpaired t-test.

## Results

### L6-silencing reduces whisker evoked multi-unit activity

As expected for WT mice, neurons in both the sensory (SCx) and motor cortex (MCx) responded consistently to low frequency (0.2 Hz) whisker pad stimulation (Table 1; Fig. 1 and 2). In the SCx, the responses consisted of both a short (peak latency <50 ms; Table 1) and long latency (peak latency >100ms) increase in firing rate. Short-latency responses represent direct ascending thalamocortical input (Moxon et al., 2008; Civillico and Contreras, 2012) and, as such, the largest responses were in L4 sensory cortex (SCx) and L5 motor cortex (MCx). Whereas the long-latency responses are mainly triggered by cortical connections (Aguilar et al., 2010; Humanes-Valera et al., 2016).

**Table 1.** Whisker evoked MUA. Firing rate (spike/s) recorded in SCx and MCx (in each layer) in control condition and 45 min after potentiation. The firing rate responses were measured in WT (SCx WT and MCx WT) and L6-Silenced animals (SCx L6-Silenced and MCx L6-Silenced). Values are mean ± SEM.

| LAYERS | Control condition | 45 min | 1 h |
| --- | --- | --- | --- |
| **SCx WT(N=9)** |  |  |  |
| **L2/3** | 7.61 ± 1.58 | 8.65 ± 1.61 | 8.37 ± 1.75 |
| **L4** | 11.84 ± 1.45 | 11.92 ± 1.50 | 12.42 ± 1.44 |
| **L5** | 10.66 ± 1.89 | 11.47 ± 1.86 | 11.64 ± 1.97 |
| **L6** | 7.14 ± 2.11 | 8.09 ± 2.16 | 8.19 ± 2.36 |
| **MCx WT(N=9)** |  |  |  |
| **L2/3** | 0.57 ± 0.22 | 1.27 ± 0.62 | 1.42 ± 0.67 |
| **L5** | 2.07 ± 0.67 | 2.27 ± 0.73 | 2.25 ± 0.67 |
| **L6** | 1.02 ± 0.76 | 0.98 ± 0.86 | 0.93 ± 0.79 |
| **SCx L6-Silenced (N=9)** |  |  |  |
| **L2/3** | 4.98 ± 0.95 <sup>#</sup> | 6.26 ± 1.62 | 6.06 ± 1.59 |
| **L4** | 8.35 ± 1.34 <sup>#</sup> | 8.90 ± 1.55 | 8.75 ± 1.48 |
| **L5** | 7.30 ± 1.25 <sup>#</sup> | 7.54 ± 1.55 | 7.64 ± 1.32 |
| **L6** | 2.82 ± 0.55 <sup>#</sup> | 3.34 ± 0.82 | 3.19 ± 0.67 |
| **MCx L6-Silenced (N=9)** |  |  |  |
| **L2/3** | 0.94 ± 0.31 | 1.14 ± 0.50 | 0.98 ± 0.42 |
| **L5** | 1.89 ± 0.72 | 1.81 ± 0.77 | 1.94 ± 0.66 |
| **L6** | 0.49 ± 0.21 | 0.50 ± 0.31 | 0.66 ± 0.33 |

<sup>#</sup> significant decrease in somatosensory cortex (SCx), Control condition: Interaction F (3, 64) = 0.1130, p = 0.9522; **WT/L6-SILENCE**, F (1, 64) = 11.07, p = 0.0015; LAYERS F (3, 64) = 5.186, p = 0.0029. Motor cortex (MCx) - Control condition: F (2, 42) = 0.3397, p = 0.7139; **WT/L6-SILENCE**, F (1, 42) = 0.06349, p = 0.8023; LAYERS F (2, 42) = 3.623, p = 0.0354

**Figure 2.**
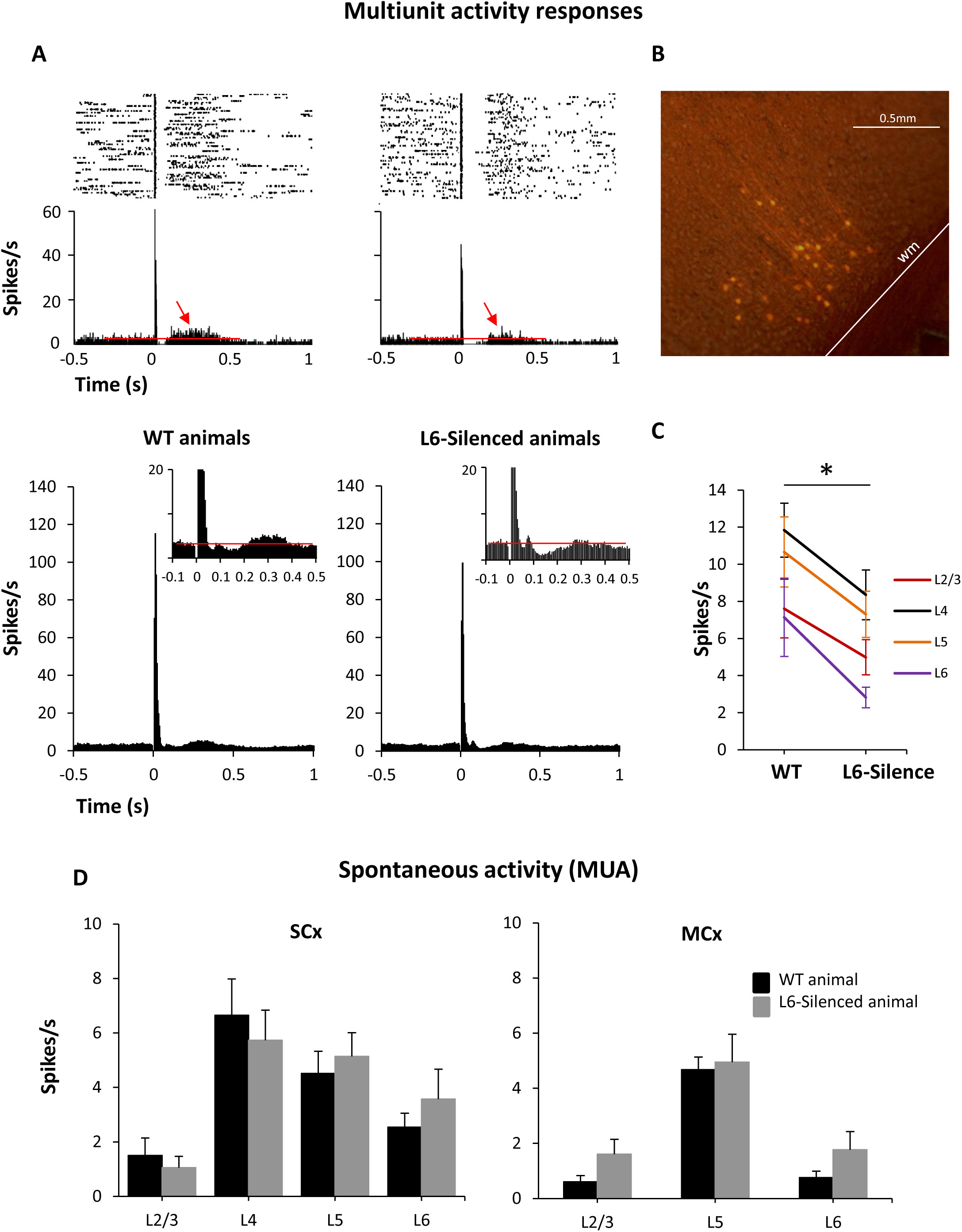
Whisker evoked multi-unit activity decreased when L6 cells were silenced. **(A)** Short and long latency whisker evoked responses. **(Top row)** Rastergrams of the MUA responses in somatosensory barrel cortex (SCx) in L4 evoked by whisker pad stimuli. Left: Example of MUA responses in L4 in a WT animal. Right: Example of MUA responses in L4 in a L6-Silenced animal. The red arrows show the long latency responses in both group of animals. **(Bottom row)** Grand average (from 9 animals) of the MUA responses evoked in SCx in L4 by whisker pad stimuli. Left: Grand averages of MUA responses in L4 in WT animals. Right: Grand averages of MUA responses in L4 in L6-Silenced animals. Inset: MUA responses truncated at 20 spikes/s to zoom in on the long latency response (latency > 100 ms). PSTH is with 5 ms bin size. **(B)** Coronal section of somatosensory cortex. Antibody staining of Ntsr1-cells (yellow) expressing tetanus toxin. The white line marks the white matter (wm). **(C)** Quantification (Table 1) of the short-latency response in each layer in WT and L6-Silenced animals shows that the whisker evoked MUA activity decreased. **(D)** Spontaneous activity (spikes per second) in different layers in SCx and MCx, in WT (black bars) and L6-Silenced (grey bars). Quantification (Table 2) plotted as a bar graph (mean ± SEM) shows that the spontaneous activity in control condition (0.2 Hz test stimulation) did not change when L6-Ntsr1 cells were blocked.

It is well established that the activity of L6-Ntsr1 cells can affect the level of cortical and thalamic excitability (Lee et al., 2012; Olsen et al., 2012; Mease et al., 2014; Reinhold et al., 2015) thus potentially influencing neuronal dynamics underlying plasticity processes. To investigate the contribution of a long-term silencing of L6-Ntsr1 pyramidal cells on sensorimotor integration, we silenced the cells by blocking synaptic release from these cells using tetanus toxin (Yamamoto et al., 2003; Nakashiba et al., 2008; Nakashiba et al., 2009). The effect of L6 silencing on cortical excitability, was investigated by measuring whisker evoked MUA activity and spontaneous activity and comparing to WT mice. Short-latency MUA responses in SCx, but not MCx, were reduced in L6-silenced animals (Table 1, Fig. 2 A,C; two-way ANOVA, SCx: WT/L6-SILENCE F(1, 64) = 11.07, p = 0.0015; MCx: WT/L6-SILENCE F(1, 42) = 0.06349, p = 0.8023), suggesting that the lack of short-latency cortical responses impacts the subsequent cortico-thalamo-cortical activity. The effect was greatest in L4 SCx, representing a 60% reduction in response magnitude, with smaller reductions of 30% in layers 2-5. However, spontaneous activity was unaffected by L6-silencing in both SCx and MCx (Table 2, Fig. 2D; two-way rm ANOVA with LAYER as within-subject main factor, SCx: WT/L6-SILENCE F(1, 19) = 0.005906, p = 0.9395; MCx: WT/L6-SILENCE, F(1, 16) = 1.896, p = 0.1874). Interestingly, the probability of a long-latency response in layer 4 was reduced after L6-silencing, being observed in 6 of 9 WT animals (67 %), but only in 3 of 9 L6-silenced animals (33 %) (Fig. 2A). Averaging over all animals the relative increased spiking in the long-latency response (ratio: number of long-latency response spikes / number of spontaneous spikes; time window 350 ms) was smaller in L6-silenced animals (unpaired t-test, p = 0.0586).

**Table 2.** Spontaneous firing rate (spike/s) recorded in SCx and MCx (in each layer) in control condition, immediately after potentiation protocol and the end of the protocol (c.f., Fig. 1). Values are mean ± SEM.

| LAYERS | Control condition | After 2Hz | End of protocol |
| --- | --- | --- | --- |
| **SCx WT (N=11)** |  |  |  |
| **L2/3** | 1.50 ± 0.65 | 1.81 ± 0.64 | 2.04 ± 0.74 |
| **L4** | 6.65 ± 1.33 | 7.91 ± 1.13 | 7.60 ± 1.12 |
| **L5** | 4.51 ± 0.81 | 5.52 ± 0.77 | 4.87 ± 0.67 |
| **L6** | 2.54 ± 0.51 | 3.41 ± 0.63 | 3.25 ± 0.61 |
| **MCx WT(N=9)** |  |  |  |
| **L2/3** | 0.61 ± 0.22 | 1.34 ± 0.53 | 1.47 ± 0.67 |
| **L5** | 4.67 ± 0.46 | 6.77 ± 0.74 | 6.72 ± 0.87 |
| **L6** | 0.76 ± 0.23 | 1.08 ± 0.29 | 1.30 ± 0.33 |
| **SCx L6-Silenced (N=10)** |  |  |  |
| **L2/3** | 1.05 ± 0.42 | 2.06 ± 0.39 | 1.86 ± 0.42 |
| **L4** | 5.73 ± 1.11 | 7.77 ± 1.25 | 7.36 ± 1.30 |
| **L5** | 5.14 ± 0.87 | 6.58 ± 1.06 | 6.31 ± 1.29 |
| **L6** | 3.57 ± 1.10 | 4.26 ± 0.82 | 3.69 ± 0.53 |
| **MCx L6-Silenced (N=9)** |  |  |  |
| **L2/3** | 1.61 ± 0.54 | 2.57 ± 1.02 | 2.39 ± 0.90 |
| **L5** | 4.95 ± 1.01 | 6.42 ± 1.06 | 7.42 ± 1.28 |
| **L6** | 1.77 ± 0.66 | 2.71 ± 0.98 | 3.30 ± 1.13 |

A whisker evoked UP state contributes to the spread of recurrent excitation across cortices (Civillico and Contreras, 2012; Wester and Contreras, 2012). The long latency responses recorded in the present study are, presumably, related (e.g., dependent on recurrent network activity) to whisker evoked UP states, thus silencing the L6 cells would reduce the propagation of activity between layers and thus reducing potentiation in the infra- and supra granular layers. In addition, our results show a reduced short latency response in every layer suggesting that the excitability of the SCx was reduced when the L6-Ntrs1 cells were silenced.

### Characterization of LFP responses in somatosensory and motor cortex

To further understand the effect of chronic L6-silencing on thalamocortical physiology, its effect on the LFP evoked response was studied. The LFP and MUA are both extracellularly recorded signals from a local network of neurons but whereas the MUA is supra-threshold activity (spikes) the LFP is mainly a reflection of synaptic inputs in the local neighborhood of the recording electrode. In WT mice, the amplitude of the LFP evoked response to whisker stimulation (0.2 Hz) was dependent on cortical layer with the largest response in L4 in SCx and L5 in MCx, respectively (Table 3, Fig. 3A,B; one-way rm ANOVA, with LAYER as within-subject factor; SCx, F(1.343, 12.09) = 12.76, p = 0.0023; MCx, F(1.605, 12.84) = 7.266, p = 0.0105; Tukey post hoc test, p < 0.05). Our results show that the cell dense L4 in SCx, and L5 in MCx receive major whisker evoked inputs. It has been shown that, for MCx, direct inputs are from both SCx, predominately from layers 2/3 and 5 (Mao et al., 2011) and sub-cortical structures (Cicirata et al., 1986; Aldes, 1988; Chakrabarti and Alloway, 2006; Feldmeyer et al., 2013). The differences between the control LFP amplitudes across the layers in both cortical areas is likely caused by a difference in subcortical and intercortical connections as others have previously observed (Di et al., 1990; Einevoll et al., 2007; Chakrabarti et al., 2008; Watanabe et al., 2014).

**Table 3.** LFP amplitude recorded in SCx and MCx (in each layer) in control condition and at different time points after 2 Hz potentiation (10, 25, 45 min and 1h). The LFP responses were measured in WT (SCx WT and MCx WT) and L6-Silenced animals (SCx L6-Silence and MCx L6-Silence). Values are mean ± SEM.

| LAYERS | Control condition | 10 min | 25 min | 45 min | 1 h |
| --- | --- | --- | --- | --- | --- |
| **SCx WT** |  |  |  |  |  |
| **L2/3** | 0.31 ± 0.07 | 0.36 ± 0.09 | 0.35 ± 0.07 | 0.39 ± 0.08 | 0.39 ± 0.08 |
| **L4** | 0.66 ± 0.14 | 0.71 ± 0.16 | 0.69 ± 0.14 | 0.74 ± 0.16 | 0.73 ± 0.14 |
| **L5** | 0.37 ± 0.07 | 0.40 ± 0.09 | 0.38 ± 0.08 | 0.42 ± 0.09 | 0.40 ± 0.09 |
| **L6** | 0.20 ± 0.03 | 0.20 ± 0.03 | 0.20 ± 0.03 | 0.20 ± 0.03 | 0.19 ± 0.03 |
| **MCx WT** |  |  |  |  |  |
| **L2/3** | 0.22 ± 0.06 | 0.27 ± 0.09 | 0.27 ± 0.09 | 0.30 ± 0.10 | 0.30 ± 0.09 |
| **L5** | 0.33 ± 0.08 | 0.39 ± 0.12 | 0.38 ± 0.10 | 0.41 ± 0.11 | 0.39 ± 0.10 |
| **L6** | 0.16 ± 0.04 | 0.20 ± 0.08 | 0.20 ± 0.06 | 0.20 ± 0.06 | 0.21 ± 0.06 |
| **SCx L6-Silenced** |  |  |  |  |  |
| **L2/3** | 0.20 ± 0.03 | 0.27 ± 0.07 | 0.25 ± 0.04 | 0.25 ± 0.04 | 0.25 ± 0.04 |
| **L4** | 0.50 ± 0.07 | 0.56 ± 0.08 | 0.56 ± 0.10 | 0.58 ± 0.10 | 0.61 ± 0.10 |
| **L5** | 0.31 ± 0.03 | 0.32 ± 0.04 | 0.33 ± 0.05 | 0.33 ± 0.05 | 0.36 ± 0.05 |
| **L6** | 0.21 ± 0.03 | 0.19 ± 0.03 | 0.20 ± 0.04 | 0.20 ± 0.04 | 0.21 ± 0.04 |
| **MCx L6-Silenced** |  |  |  |  |  |
| **L2/3** | 0.17 ± 0.04 | 0.17 ± 0.03 | 0.17 ± 0.04 | 0.20 ± 0.05 | 0.21 ± 0.04 |
| **L5** | 0.29 ± 0.07 | 0.28 ± 0.06 | 0.28 ± 0.08 | 0.31 ± 0.08 | 0.33 ± 0.08 |
| **L6** | 0.14 ± 0.02 | 0.14 ± 0.02 | 0.12 ± 0.01 | 0.15 ± 0.02 | 0.15 ± 0.02 |

SCx WT. Two-way rm ANOVA: Interaction $F(12,108) = 2.796$ , $p = 0.0023$ ; LAYERS $F(3,27) = 13.83$ , $p < 0.0001$ ; TIME $F(4, 36) = 2.607$ , $p = 0.0518$ . (N=10)
MCx WT. Two-way rm-ANOVA: Interaction $F(8,64) = 1.060$ , $p = 0.4024$ ; LAYERS $F(2,16) = 4.641$ , $p = 0.0257$ ; TIME $F(4,32) = 2.816$ , $p = 0.0414$ (N=9)
SCx L6-Silence. Two-way rm-ANOVA: Interaction $F(12,108) = 1.567$ , $p = 0.1120$ ; LAYERS $F(3,27)=15.74$ , $p<0.0001$ ; TIME $F(4,36) = 1.492$ , $p = 0.2250$ (N=10)
MCx L6-Silence. Two-way rm-ANOVA: Interaction $F(8,64) = 0.7708$ , $p = 0.6296$ ; LAYERS $F(2,16) = 5.549$ , $p = 0.0148$ ; TIME $F(4,32) = 1.326$ , $p = 0.2814$ (N=9)

**Figure 3.**
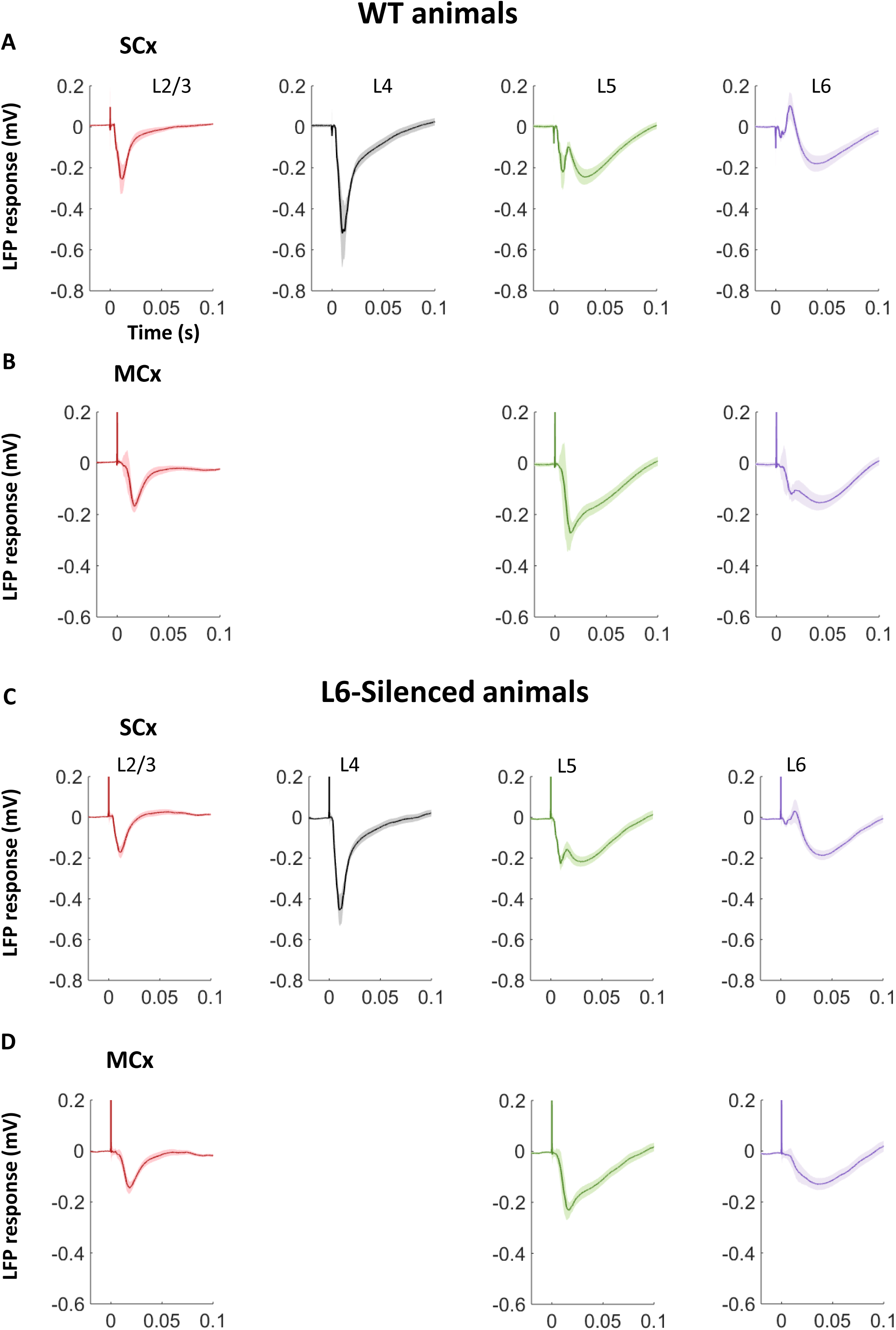
Characterization of LFP responses in somatosensory and motor cortex. **(A-B)** The average LFP response in SCx and MCx, evoked by electrical stimuli delivered to the whisker pad in WT animals (Table 3). **(A)** Grand average of LFP responses in the different SCx layers and SEM. (B) Grand average of LFP responses in the different MCx layers. **(D-E)** The average of LFP response in SCx and MCx by electrical stimuli delivered to the whisker pad in L6-Silenced animals (Table 3). **(D)** Grand average of LFP responses in the different SCx layers. **(E)** Grand average of LFP responses in the different MCx layers. All data plotted as mean ± SEM.

Silencing L6 activity did not affect the amplitude of the evoked response in sensory or motor cortex (Table 3, Fig. 3; two-way rm ANOVA with LAYER as within-subject main factor; SCx: WT/L6-SILENCE, F(1, 18) = 1.003, p = 0.3298; MCx: WT/L6-SILENCE, F(1, 16) = 0.2693, p = 0.6109), suggesting that thalamocortical activity was not affected. Taken together, these data suggest that L6 modulates cortical responsiveness without affecting resting state activity or synaptic input.

### L6-silencing modulated synaptic plasticity

These effects of L6-silencing on cortical excitability could influence neuronal dynamics underlying plasticity processes. To study this, the whisker pad was stimulated electrically at 2 Hz for 10 min to produce plasticity in the somatosensory and motor cortex, as previously shown (Megevand et al., 2009; An et al., 2012); see Fig. 1A for the stimulation protocol). In WT mice the whisker pad stimulation protocol potentiated the LFP in both SCx and MCx (Table 3; Fig. 4). The effect was layer specific such that in SCx the evoked LFP amplitude after 45 min increased by 25 % in L2/3 and by 12% in L4 (Table 3, Fig. 4A), but with no significant effect in L5 and L6 (Table 3; two-way rm ANOVA matching both main factors). The increase in the LFP amplitude reached significance 10 min after the potentiation stimulation (Dunnett’s test; L2/3: p = 0.0069; L4: p = 0.0046), and remained potentiated during the one-hour measurement (Dunnett’s test, p<0.05, and one-way rm ANOVA post-test for linear trend L2/3: p = 0.0002; L4 p = 0.0116). Similarly, in MCx, L2/3 and L5 was potentiated (Table 3, Fig. 4B; 36% in L2/3: 24% in L5) within 10 min of the start of stimulation (Dunnett’s test; L2/3: p = 0.0107; L5: p = 0.0006) and remained potentiated for at least one hour (Dunnett’s test, p<0.05, and one-way rm ANOVA post-test for linear trend L2/3: p = 0.0015, L5: p = 0.0488). These findings show a layer-specific potentiation in both cortical areas, with the potentiation in somatosensory cortical L2/3 presumably contributing to the spread of the potentiation to motor cortex.

**Figure 4.**
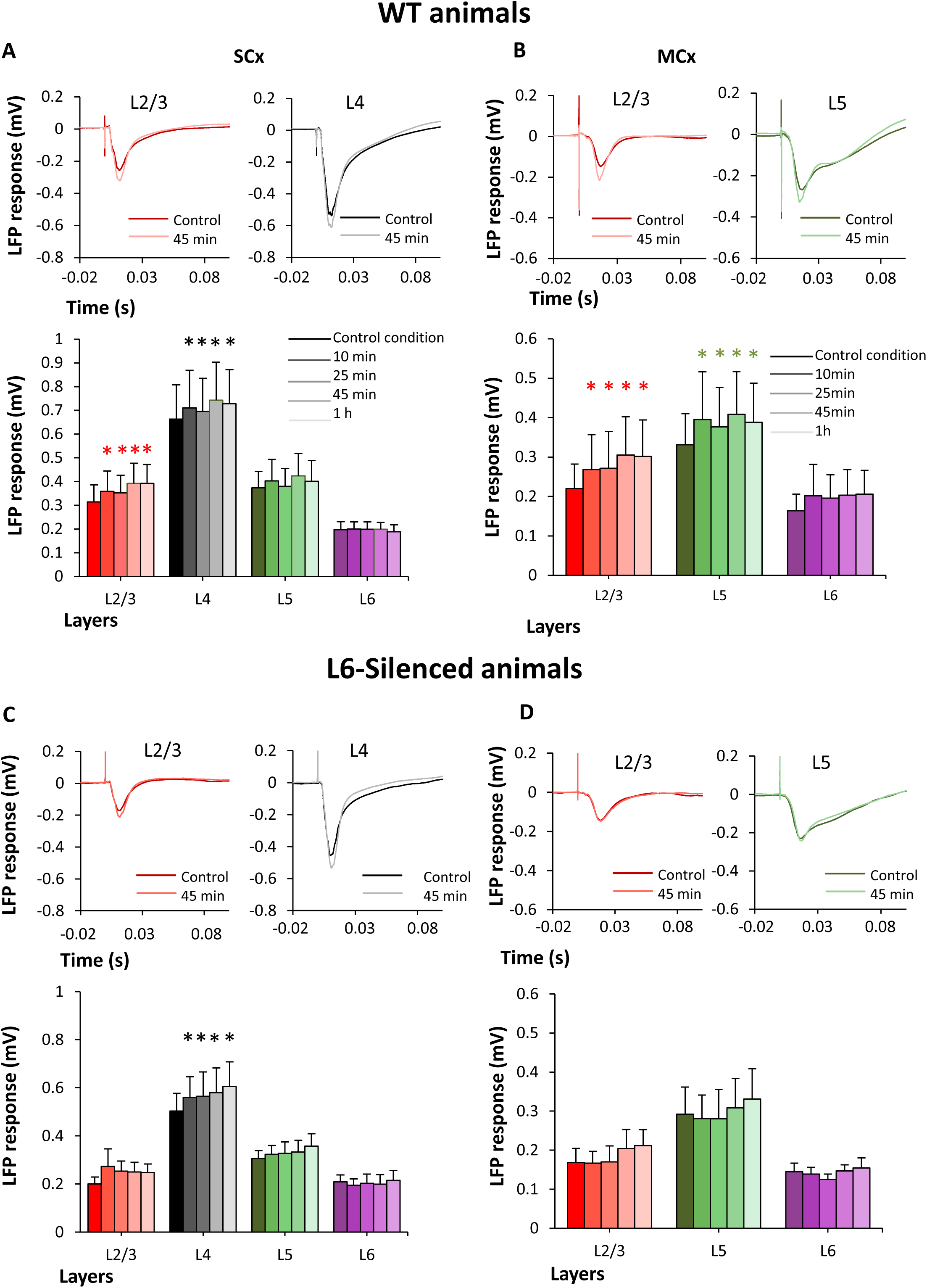
L6 corticothalamic cells contribute to the propagation of potentiation. (**A-B**) Traces show average LFP responses, and histograms show summary the efficacy of the stimulation protocol to induce potentiation in different layers in WT animals. (**A**) *Top row*: Grand average of LFP responses in L2/3 and L4 in SCx in control condition and 45 min after the potentiation protocol. *Bottom row*; Quantification of the average LFP response over time in SCx. (**B**) *Top row*: Grand average of LFP responses in the L2/3 and L5 in MCx in control condition and 45 min after the potentiation protocol. *Bottom row*: Quantification of the average LFP response over time in MCx. (**C-D**) Traces show average LFP responses, and histogram show the efficacy of the stimulation protocol to induce potentiation in different layers in L6-Silenced animals. (**C**) *Top row*: Grand average of LFP responses in the L2/3 and L4 in SCx in control condition and 45 min after potentiation protocol. *Bottom row*: Quantification of the average LFP response over time in SCx. (**D**) *Top row*: Grand average of LFP responses in the L2/3 and L5 in MCx in control condition and 45 min after potentiation protocol. *Bottom row*: Quantification of the average LFP response over time in MCx. Stars mark a significant difference (2-way ANOVA with Dunnett’s test) to control conditions within each respective layer. Numbers to the bar graphs in Table 3. Mean ± SEM.

We next examined the effect on the short-latency MUA evoked response. The potentiating protocol did not alter the short latency evoked MUA in SCx nor in MCx (Table 2: two-way rm ANOVA, matching both main factors. TIME: WT-SCx: F(2, 16) = 1.153, p = 0.3407; WT-MCx: p = 0.2114), nor did it affect the spontaneous MUA in SCx (Table 2; two-way rm ANOVA matching both main factors. TIME: (i) SCx-WT: F(2, 20) = 2.367, p = 0.1195). However, it did affect spontaneous MUA in MCx. In motor cortex, the spontaneous MUA increased over time (Table 2; two-way rm ANOVA matching both main factors, MCx-WT: Interaction F(4, 32) = 2.678, p = 0.0494, TIME F(2, 16) = 4.672, p = 0.0252, LAYER F(2, 16) = 100.7, p < 0.0001).

With a better understanding of how the potentiating protocol modulates the responsiveness of SCx and MCx to sensory stimulation, we next examined the impact of silencing L6 pyramidal cells on the potentiation of LFP evoked response in both cortical regions.

Silencing the output of L6-Ntsr1 cells in SCx using tetanus toxin we observed a blocked potentiation of the LFP response (Fig. 4C,D) in both SCx and MCx (Table 3; two-way rm ANOVA matching both main factors (LAYER AND TIME). The lack of an interaction effect in SCx L6-Silenced would imply that potentiation is completely blocked in both layers 2/3 and 4. A post hoc analysis suggests, however, that the potentiation is not completely blocked in L4 (Dunnett’s test, p < 0.05 at all 4 time points, and one-way rm ANOVA significant post test for linear trend in L4 (p = 0.0106) but not L2/3 (p = 0.3921)). Since L6 does not affect synaptic inputs to SCx (see L6-silencing Does not affect LFP Evoked Responses, above), L6 likely modulates long-term plasticity through mechanisms at the cortical level.

To better identify the likely cortical mechanisms, we examined the effect of L6-silencing on spontaneous and evoked MUA activity since they have been shown to affect the initiation of potentiation (Arieli et al., 1996; Azouz and Gray, 1999; Tsodyks et al., 1999; Lakatos et al., 2008). Interestingly, spontaneous MUA in SCx was not affected by the potentiation protocol in L6-silenced animals (Table 2; two-way rm ANOVA matching both main factors. (TIME: (ii) SCx-SILENCE: F(2, 18) = 2.582, p = 0.1033) and spontaneous MUA in MCx increased over time (Table 2; two-way rm ANOVA matching both main factors, MCx-L6-SILENCE: Interaction F(4, 32) = 1.160, p = 0.3468, TIME F(2, 16) = 3.911, p = 0.0414, LAYER F(2, 16) = 7.625, p = 0.0047). This increase in spontaneous MUA in MCx, but not SCx is similar to the effect in WT noted above. Furthermore, the potentiating protocol under L6-silencing did not affect the evoked MUA in SCx nor in MCx (L6-SILENCE-SCx: F(2, 16) = 1.613, p = 0.2301; L6-SILENCE-MCx: p = 0.9094), again, similar to WT MUA evoked responses. The increase in spontaneous MUA in MCx, but not SCx suggests that sub-cortical pathways projecting to motor cortex could be affected independently of neocortex.

### Effect of silencing L6 cells on behavior in a whisker-dependent behavioural task

Since L6-silencing modulates long-term plasticity, it is important to understand the behavioral consequences. Animals were tested in a tactile-dependent behavioral task designed to investigate sensory-motor learning abilities (Hutson and Masterton, 1986; Harris et al., 1999; Celikel and Sakmann, 2007; Papaioannou et al., 2013; Juczewski et al., 2016) where they were required to reach across a gap to a platform, use their whiskers to determine the width of the gap, and then jump to the platform. WT and L6-Block animals both performed this task equally well judging from the fact that the max gap distance reached was 5.25 cm for both groups (averaged over the two last days unpaired t-test p = 1) and they had the same percent of successful attempts (WT: n = 7, 9.3 ± 1.5 %; L6-Silenced: n = 10, 11.9 ± 1.9%; unpaired t-test, p = 0.33). Furthermore, we did not observe a significant difference in time the animals spent exploring the platform “Exploration Duration Successful Attempt” (WT: n=9, 3.67±0.3 s; L6-Silenced: n=10, 3.05±0.31 s; unpaired t-test, t=1.391 df=15, p=0.18) and the duration of successful attempts (WT: n=7, 13.5 ± 1.2 s; L6-Silenced: n = 10, 11.6 ± 1.0 s; unpaired t-test, t=1.207 df=15, p = 0.25) and unsuccessful (WT: n=9, 4.0 ± 0.5 s; L6-Silenced: n = 10, 3.9 ± 0.4 s; unpaired t-test, t=0.2065, df=17, p = 0.84). To investigate how silencing L6 cells modifies the acquisition of the tactile sensory information via the whiskers, we analyzed whisker kinematics. The data was divided into two categories: (1) short gap-distances (4.0 to 5.0 cm), where the distance is short enough that the animals can also use their nose to locate the platform thus any defects in processing of whisker information can be compensated for, and (2) long gap-distances (5.5 to 7.5 cm) where the animals can only reach the platform with its whiskers, thus any changes in the go/no-go decision based on whisker input is more prominent. Whisker-dependent sensory perception was quantified by measuring whisker contacts with the platform and whisker amplitude. As expected for the short gap-distances, WT and L6-Silenced animals did not show any difference in the number of whisker contacts in successful attempts (WT: 9.8 ±0.6; L6-Silence: 8.7 ± 0.6; unpaired t-test, t=1.221, df=15, p = 0.24) or the duration of each contact (WT: 35.0 ± 1.4 ms; L6-Silence: 31.4 ± 1.7 ms; unpaired t-test, t=1.506, df=15, p = 0.15). In contrast, at longer distances the L6-Silence animals made fewer whisker contacts (Fig. 5A, WT: 15.1 ± 1.5; L6-Silence: 9.7 ± 1.5; unpaired t-test, t=2.401, df=10, p = 0.037). In addition, we observed that the duration of the whisker touch was shorter in L6-Silence animals (Fig. 5B; WT: 36.2 ±1.6 ms; L6-Silence: 30.5 ± 1.7 ms; unpaired t-test, t=2.365, df=10, p = 0.039). Furthermore, he protraction amplitude was measured and a noticeable difference was that in L6-Silenced animals there was relatively more of low amplitude (amplitudes < 20 deg) whisks (Fig. 5C,D,E, Kolmogorov-Smirnov independent sample test, p<0.001; chi-square test p < 0.0001).

**Figure 5.**
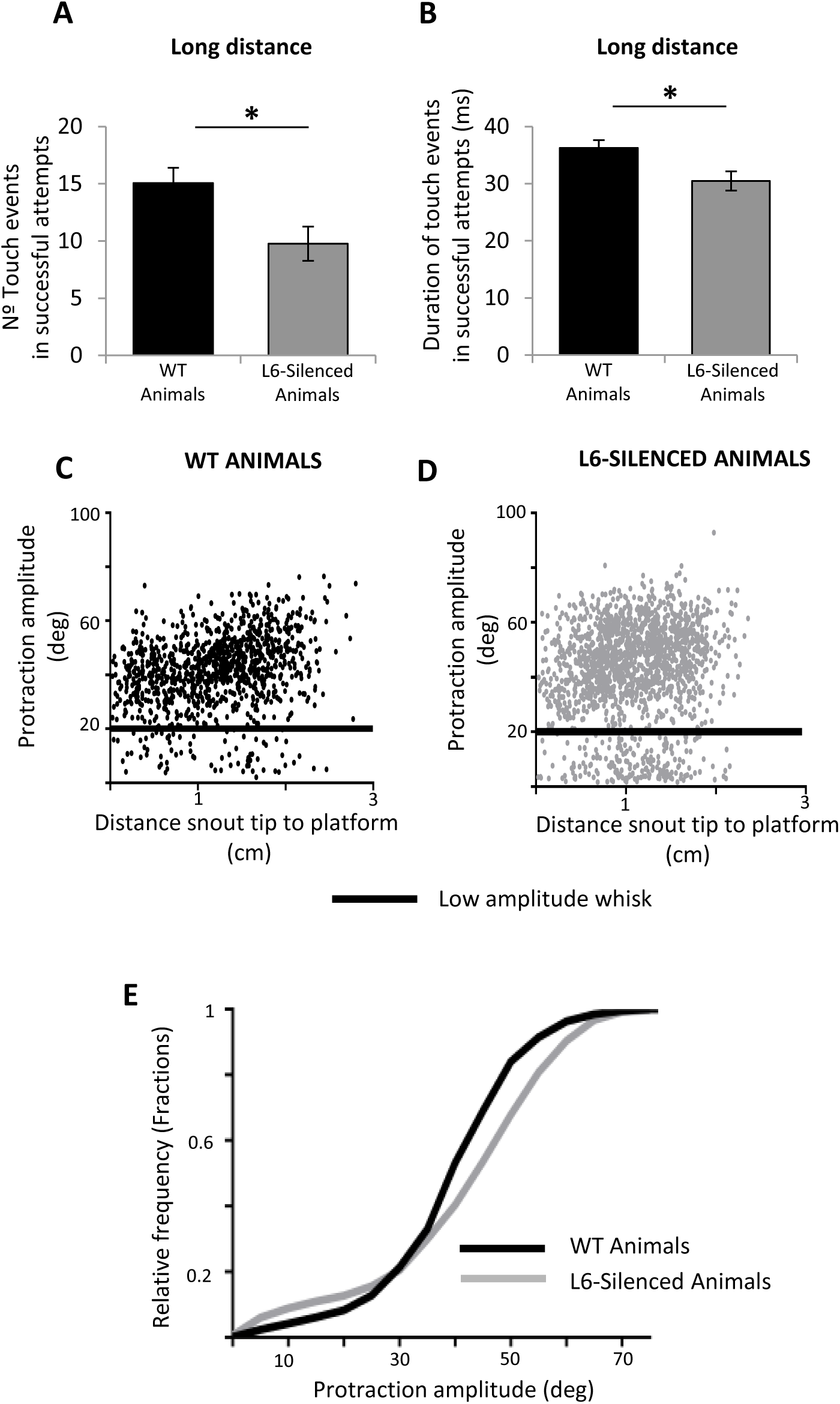
Effect of silencing L6 cells on behavior in a whisker-dependent behavioural task. **(A-B)** Histograms show parameters used to quantify the animals use of its whiskers to explore the target platform for long gap-distances (5.5 – 7.5 cm). **(A)** The number of touch events in successful gap-crossing attempts. Mean ± SEM**. (B)** The duration of touch events in successful gap-crossing attempts. * p < 0.05. **(C-D)** Scatter plot shows properties of whisker kinematics in WT and L6 silenced animals. **(C)** Whisker amplitudes in WT animals (y-axis) as a function of snout tip distance (in cm) to the platform. **(D)** Whisker amplitudes in L6-Silenced animals. (y-axis) as a function of snout tip distance (in cm) to the platform. Black line added as reference for “low amplitude” whisks with an amplitude < 20 deg. (E) Cumulative frequency distribution of whisker amplitudes in WT (Black curve) and in L6-Silenced animals (Grey curve) show that there are relatively more low amplitude whisks in the L6-Silenced animals (KS-test, p p<0.001).

## Discussion

The present study shows the importance of L6-Ntsr1 pyramidal cells in regulating cortical plasticity. Permanent silencing of the L6-Ntsr1 cells reduced the whisker evoked MUA responses across all layers of SCx with the greatest impact on L6. Importantly, high frequency stimulation induced potentiation of the LFP response in SCx and MCx, and blocking L6 activity reduced the potentiation in both cortical areas. Chronical silencing of L6 activity thus reduced cortical excitability, and cortical plasticity which has impact on processing of sensory information. In the behaving animal this reduction appears to be compensated for such that the animal uses low-amplitude whisks (Towal and Hartmann, 2008) when performing a tactile detection task.

### L6-silencing reduces whisker evoked multi-unit activity

To study the efficacy of silencing of L6-Ntsr1 cell and to analyze the L6 contribution to sensory processing we analyzed the multi-unit activity (MUA) from somatosensory and motor cortex. We observed that MUA “responses” were affected by blocking L6 (Fig. 2A). Short-latency responses, typical of somatosensory responses (Moxon et al., 2008), and long-latency responses, which are triggered UP states (Hasenstaub et al., 2007; Reig and Sanchez-Vives, 2007; Humanes-Valera et al., 2016), were reduced in L6-Silenced animals (Fig. 2A,C). L6 cells can modulate the cortical activity by local connections in the same layer and with other layers in the same area of the cortex (Lefort et al., 2009). L6 cells have strong connections with inhibitory interneurons (Bortone et al., 2014; Kim et al., 2014). Different types of interneurons are involved in the modulation of UP states (Puig et al., 2008; Neske and Connors, 2016) therefore silencing L6 cells we are presumably decreasing the activity of interneurons, thus causing a reduction in the generation of UP states associated with the sensory stimuli. The propagated UP state is necessary for the spread of recurrent excitation across cortices (Civillico and Contreras, 2012; Wester and Contreras, 2012) consequently these results suggest that when L6 is blocked whisker evoked active states are reduced possibly reflecting a decreased activation and capacity to propagate cortical UP states. Silencing the L6 over an extended period did not affect the spontaneous activity (Fig. 2D), whisker evoked responses were, however, reduced when L6 activity was chronically blocked (Fig. 2A). Apart from differences in the experimental protocol, one explanation could be in how L6 modulates global cortical activity suggesting that plasticity mechanisms cause cortical changes during the time of tetanus toxin incubation.

### Characterization of LFP responses in somatosensory and motor cortex

The LFP is a local measure of both sub-threshold synaptic activity and spiking (Mitzdorf, 1985; Logothetis et al., 2001; Einevoll et al., 2007; Nielsen and Rainer, 2008; Katzner et al., 2009; Linden et al., 2011; Watanabe et al., 2014). Our results show a difference in the LFP amplitude between layers in both cortical areas (Fig. 3A,B) that is presumably caused by a difference in subcortical and inter-cortical connections as others have previously observed (Di et al., 1990; Einevoll et al., 2007; Chakrabarti et al., 2008; Watanabe et al., 2014).

Analyzing changes in the LFP is a useful measure to study how activity spreads between layers and cortices (Kleinfeld et al., 2006; Chakrabarti et al., 2008; Hooks et al., 2011; Zagha et al., 2013; Petrof et al., 2015; Hooks, 2016). The LFP amplitude in each respective SCx layer was not significantly different between WT and L6-Silenced animals (Table 3, Fig. 3) indicating that an effect on the input from thalamus was not the major cause underlying the reduced potentiation when L6 cells were silenced.

### L6-silencing modulated synaptic plasticity

Rodents explore the environment moving their whiskers at different frequencies depending on behavioral context (Carvell and Simons, 1990; Wolfe et al., 2008). In the cerebral cortex, sensory stimuli at certain frequencies have been described to produce synaptic plasticity such as LTP (Clapp et al., 2005; Troncoso et al., 2007; An et al., 2012; Han et al., 2015). We induced a potentiation in somatosensory cortex using a 2 Hz stimulation protocol (Megevand et al., 2009; An et al., 2012). The potentiation was induced in SCx L2/3 and L4, but no detectable potentiation was found in L5 and L6 (Fig. 4A). A similar layer specificity with the strongest potentiation in layers 2-4 was also reported for passive whisker stimulation in rats (Megevand et al., 2009; An et al., 2012). The increased whisker evoked LFP amplitude in L4 of SCx suggests that activity along the lemniscal pathway was potentiated by the peripheral whisker pad stimulation (Lee and Ebner, 1992). Layer 4 cells project mainly to L2/3 (Radnikow et al., 2015) and thus contribute to cortical plasticity occurring in L2/3 (Allen et al., 2003; Shepherd et al., 2003; Bender et al., 2006; Clem and Barth, 2006; Hardingham et al., 2008; Krieger, 2009; House et al., 2011; Gambino et al., 2014; Ma et al., 2016). Direct lemniscal inputs to cortex is distributed in both layers 4 and 5B (Constantinople and Bruno, 2013), with L5A also receiving extensive inputs from the paralemniscal pathway (Ahissar et al., 2001; Groh et al., 2010; Oberlaender et al., 2012; Feldmeyer et al., 2013). In our experimental paradigm the LFP amplitude in L4, but not in L5, of SCx was potentiated, suggesting that the whisker stimulation primarily potentiated activity via the lemniscal pathway. The direct connection between SCx and MCx is important for sensorimotor integration (Mao et al., 2011; Petrof et al., 2015), and axons from L2/3 and L5 in SCx project to L2/3 and L5 in vibrissal MCx (Hooks et al., 2011; Mao et al., 2011; Petrof et al., 2015). Our results are in agreement with these observations because potentiation was observed in L2/3 in SCx that was then presumably propagated to L5 and L2/3 in MCx (Fig. 4).

Silencing the output of the L6-Ntsr1 cells, the potentiation of the LFP response was affected in both cortical areas. In SCx of L6-Silenced animals, there is some evidence in support of a small potentiation remaining in L4, but not in L2/3. This could indicate that the thalamic connection to L4 in SCx was still potentiated, but to a lesser degree, and this smaller potentiation was not sufficient for the potentiation to spread to L2/3 (Fig. 4C). One explanation of this is the reduction of propagated UP states required to spread recurrent excitation across cortices (Civillico and Contreras, 2012; Wester and Contreras, 2012). These results suggest that when L6 is blocked, triggered UP states are reduced, possibly reflecting a decreased level of activation and capacity to propagate cortical UP states. It is intriguing to consider that L6 cells are involved in cortical plasticity facilitating the triggered UP states to spread across the layers. In addition, the potentiation in MCx was also affected (Fig. 4D). Since the reduction in SCx was in L2/3, which projects to MCx, it can be assumed that this reduction contributes to a reduced potentiation in MCx. This is in agreement with MCx being less prone to synaptic plasticity (Castro-Alamancos et al., 1995). Since MCx also receives direct sub-cortical projections (Asanuma and Hunsperger, 1975; Porter and White, 1983; Hooks et al., 2013) these connections could also influence potentiation in MCx. If the thalamo-cortical synapse remains potentiated when the L6-Ntsr1 cells are blocked, this could indicate that L6, depending on the sensory input parameters, can modulate cortical and thalamic activity to different degrees (Briggs and Usrey, 2008).

One explanation to a reduction of the sensory excitability could be an excessive increase in spontaneous activity (Aguilar et al., 2010; Humanes-Valera et al., 2013; Yague et al., 2014; Alonso-Calvino et al., 2016; Humanes-Valera et al., 2016). The spontaneous activity was recorded before and after the stimulation protocol (see Fig. 1), and we conclude that the level of spontaneous activity was not changed in the L6-Silenced animals. The loss of potentiation of evoked LFP after silencing L6 is not simply due to a reduction in general excitability but likely alters some specific mechanisms of long-term plasticity.

### Effect of silencing L6 cells on behavior in a whisker-dependent behavioural task

To understand the behavioral consequences of permanently silencing L6 cells, we tested the L6-block animals in a tactile-dependent task designed to investigate sensory-motor learning abilities (Hutson and Masterton, 1986; Harris et al., 1999; Celikel and Sakmann, 2007; Papaioannou et al., 2013; Juczewski et al., 2016). Somatosensory cortex responds with bigger responses preferentially to high-velocity inputs producing more synchronization in sensory thalamus (Pinto et al., 1996; Pinto et al., 2000; Temereanca and Simons, 2003; Towal and Hartmann, 2008). The L6-block animals have reduced MUA responses in the SCx, to compensate this reduction, the animal adds low-amplitude whisks to the normal whisking cycle. This mechanism could thus potentially prime the system to achieve an improved information acquisition (Fig. 5C,D). The reduction of the number of touches used to solve the task could be due to the compensatory strategy with brief low-amplitude whisks that made the system more efficient (Fig. 5A,B).

The dual projections of L6 cells to both cortex and thalamus enables them to control both a “feed-back” circuit to thalamus, but also their effect on cortical activity can be seen as a “feed-forward” modulation. These terms seems to imply a serial processing route, where we can actually establish “who’s on first and what’s on second”, and should perhaps only be used to emphasis that the corticothalamic cells have different projection with different tasks, rather than suggesting a strict information processing pathway (Thomson, 2010). Corticothalamic L6 pyramidal cells have an important function in the thalamo-cortico-thalamic circuit crucial for animal perception. In this work, we investigated the contribution of these cells in somatosensory cortex to cortical plasticity and sensorimotor connections. Silencing a sub-population of corticothalamic L6 cells, the whisker evoked responses were reduced in somatosensory cortex. During a potentiation protocol used to study a cortical plasticity, the LFP potentiation in both cortices was affected. The neuroplasticity induced by chronic silencing of Layer 6 is compensated behaviorally by modulating sensory acquisition (whisking) to compensate for the loss in excitability.

## Author contribution

D.H-V and P.K designed research; D.H-V performed extracellular recordings with silicon probes; V.G. did the behavioral experiments and analyzed the data; J.A and K.M developed analysis tools and M.S viral constructs. D.H-V and P.K analyzed data and wrote the paper.

## Acknowledgements

We thank Ann-Christin Ammann for technical assistance, and Juan Aguilar and Guglielmo Foffani for comments on an earlier version of the manuscript. This project was supported by the Deutsche Forschungsgemeinschaft (DFG) grant SFB 874/A9 to P.K.

## Ethical approval

All applicable international, national, and/or institutional guidelines for the care and use of animals were followed.

